# Nucl2Vec : Local alignment of DNA sequences using Distributed Vector Representation

**DOI:** 10.1101/401851

**Authors:** Prakhar Ganesh, Gaurav Gupta, Shubhi Saini, Kolin Paul

## Abstract

The Next Generation Sequencing Technique (NGS) has provided affordable and fast method for generating genetic data. Generation of whole Genome Sequence and extract relevant information from this data is still a computationally expensive process. In this paper we demonstrate a novel approach for local alignment of DNA reads with respect to reference genome. For this process we have used Skip-gram model for creating encoding(Nucl2Vec) and k-nearest neighbor for the alignment. With our new approach we have reduced computation cost for local alignment, while achieving accuracy comparable to existing defacto standard BWA-MEM tool.

**Index Terms:** Genome, Alignment, Local Alignment, k-nearest Neighbor, Distributed vector representation, Skip-gram

## I. Introduction

Genome sequencing is figuring out the order of DNA nucleotides(bases) Adenine, Guanine, Cytosine and Thymine (A,C,G and T respectively) that make up an organism’s DNA. The whole genome cannot be sequenced all at once because available methods of DNA sequencing can only handle short stretches of DNA at a time [1]. Therefore, the genome is sequenced in pieces (whole-genome shotgun method). In this method, the genome is broken up into small pieces which are then sequenced. To regenerate whole genome sequence, these reads are either assembled using existing reference DNA(alignment) or by using de-novo assembly [2]. The reads generated using High Throughput sequencer are sequenced randomly and are of fragment length of 100-900 bps with error rate of 1%-2% per read [3]. To facilitate variant discovery with high confidence these reads are generated with average 30x coverage. Due to large number of short length reads produced with high sampling rate, their alignment, along with error isolation is a computationally intensive task.

Alignment is the process by which we align the NGS reads to their corresponding(most likely) locations in the reference [4]. Computational approaches to sequence alignment generally fall into two categories: global alignments and local alignments. Calculating a global alignment is a form of global optimization that forces the alignment to span the entire length of all query sequences. By contrast, local alignments identify regions of similarity within long sequences that are often widely divergent overall. (Fig. 1)

**Fig. 1.**
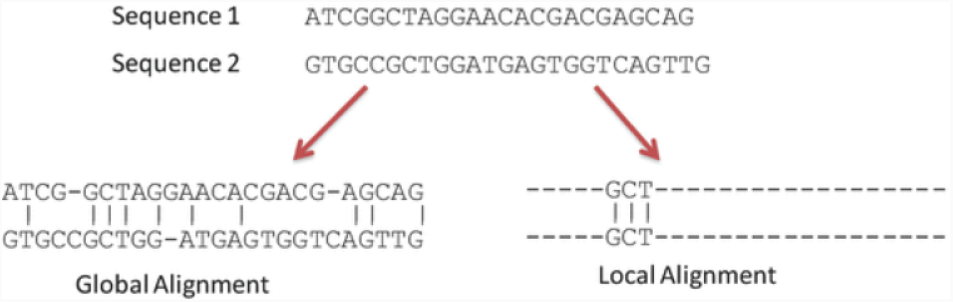
Global vs Local Alignment

## II. Existing Technique

To generate Whole genome sequence from raw NGS reads, BWA-MEM and BLAST are two of the most popular tools. BWA-MEM is based on Burrows-Wheeler Transform algorithm and follows the canonical seed-and-extend paradigm which finds at each query position the longest exact match covering the position [5]. Whereas Basic Local Alignment Search Tool(BLAST) is based on local alignment of read against reference genome. It is a heuristic search which seeks word of Length W that score at least T when aligned with the query. Words in the database that score T or greater are extended in both directions in an attempt to find a locally optimal ungapped alignment known as high scoring pair(HSP). [6]. Both BWA and BLAST algorithms use dynamic programming, which solves a problem by dividing problem into smaller problem and then growing solution from subproblems. There have been other techniques such as suffix trees [7], KMS algorithm [8], fuzzy logic [9] and greedy method [10] that are faster than dynamic programming.

In this paper we are proposing a Machine Leaning Algorithm using K- nearest neighbor to align DNA Reads w.r.t reference genome. Any machine learning based model for alignment of reads can be broadly divided into two parts: (a) feature engineering and (b) applying the machine learning algorithm. The features extracted from raw data are fed as input to the machine learning algorithm. This makes feature extraction a critical step in the success of any ML algorithm. While some work has been done for identification and development of algorithms suitable for alignment [11] [12], little has been done in extracting feature information from DNA.

Information in DNA is represented in the form of sequence of 4 characters, as described earlier. Thus, we require a suitable model to encode this string sequence into a mathematical representation. The most commonly used encoding is one-hot vector encoding, which represents each nucleotide as a vector of size 4. However, this encoding does not translate any sequence related information (bi-nucleotide, tri-nucleotide relationships etc.), moreover it increases the dimension by four folds and this degrades the efficiency of ML algorithm. Providing better encodings translates compressing more relevant information into lower dimension which can boost the performance of any machine learning algorithm that we use. Since a set of nucleotides in DNA sequence can be represented as word, we propose an encoding similar to that of Word2Vec[2, 3] to encode sequential information of k-mers before feeding into a machine learning algorithm. We have named this encoding as Nucl2Vec encoding.

## III. Proposed Solution

Our proposed solution is based on generating Nucl2Vec encodings of the input reference and read sequence k-mers. The reference k-mer encodings are used to generate a K Nearest Neighbors(KNN) tree. This is a one time activity. Subsequently, for each read, predicted position is obtained through a search in this KNN tree based on its encoding. (Fig 3)

**Fig. 3.**
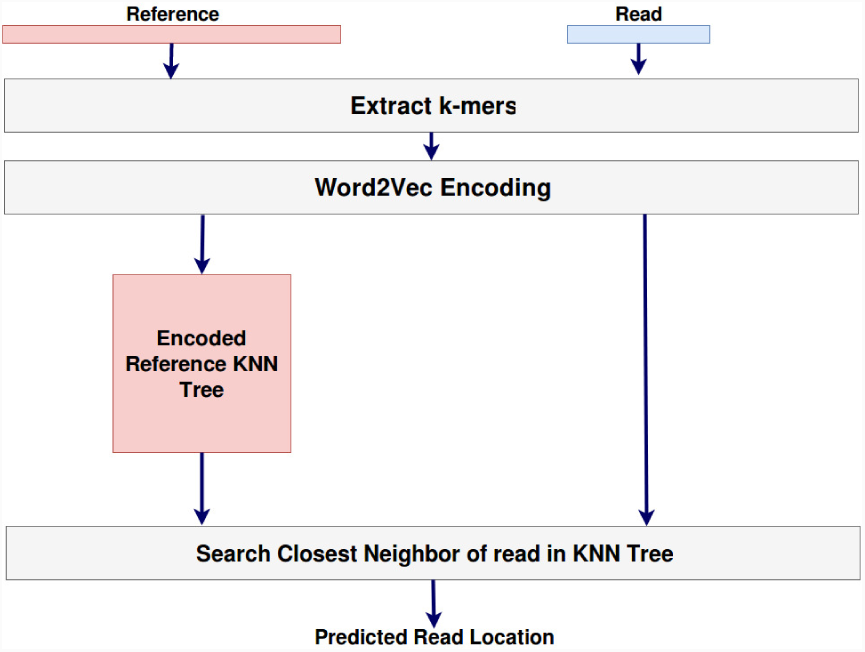
Model Overview

For alignment, instead of considering the whole read sequence, we use only a portion of the read (see Fig 2) to create encodings. By using this heuristic, we are improving the performance of KNN by reducing the size of search space. This is based on the intuition of how we solve jigsaw puzzles. While joining two pieces of a jigsaw puzzle, we only consider the corners and not the content present in the center. Similarly, for finding correct alignments for the reads, we only consider small portions of the read and hypothesize that the rest of the read will match at the same place where the smaller portion matches. Following subsections describe encoding of DNA data and use of k-nearest Neighbor for local alignment.

**Fig. 2.**
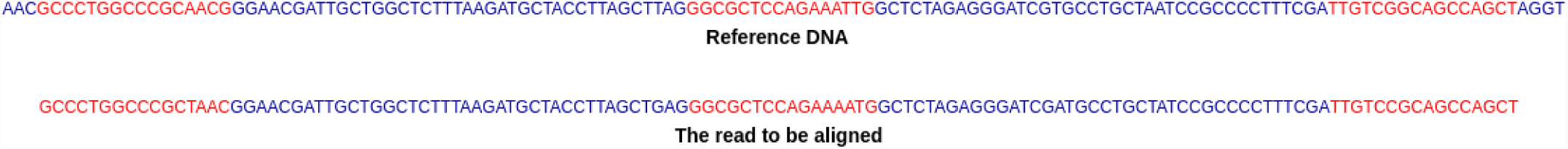
Matching only a portion of the read (represented by red). The rest of the read (represented by blue) seems to match at the same location too.

### A. Nucl2Vec Encoding

Word2Vec has been typically used as a feature engineering technique for Natural Language Processing related problems. However, we aim to introduce this concept to our encoding of DNA base pair sequence. Word2Vec, based on the Skip-gram model, is an efficient method for learning high quality vector representations of words from large amounts of unstructured text data. Unlike most of the previously used neural network architectures for learning word vectors, training of the Skip-gram model does not involve dense matrix multiplications. This makes the training extremely efficient [13].

Word2Vec in NLP is trained using heuristics like negative sampling, subsampling and Dynamic context windows [14]. This is required due to the huge size of vocabulary in Natural Language and sparseness of correlations between the words. However in our case, since our vocabulary was smaller in size we forgo these heuristics and stick to hierarchical softmax without subsampling. Though to encode our k-mers into a very low dimension required more thorough training, but due to small vocabulary, the training time was not adversely affected by these choices.

Word2Vec uses proximity of words in a sentence for similarity between words. Since our vocabulary has k-mers from NGS reads, these reads may differ from reference due to substitution and indels. To accommodate this property, in Nucl2Vec, we consider the number of edits to measure degree of similarity. An edit between two k-mers can be defined as one of the following(refer Fig 4:

**Fig. 4.**
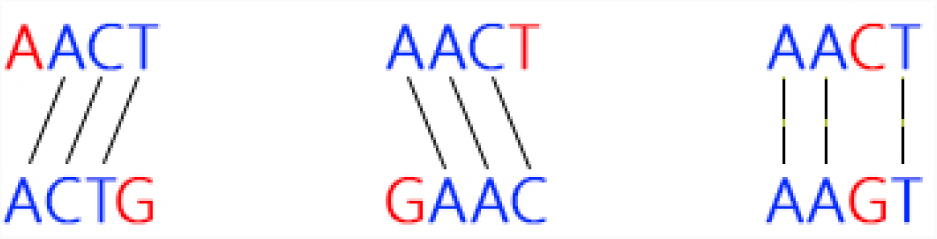
Types of Edits

1. Add a base pair on the left and shift the k-mer to the right
2. Add a base pair on the right and shift the k-mer to the left
3. Substitute a base pair

*Training Nucl2Vec Model for DNA* : We train our Nucl2Vec model to learn the edits based similarity between various k-

**Algorithm 1** Nucl2Vec Traning

~~~
**function** NEARBYSET(inputKmer) *⊳* Function to generate all k-mers at an edit distance ‘1’ from the inputKmer
  **for** every location lo in the inputKmer **do**
        *b ←* [’A’, ’C’, ’T’, ’G’]
        **for** every base pair b **do**
              *substituteKmer←*Substitution at location lo with base pair b
              *insertionKmerRight←*Insertion at location lo with base pair b and shift right
              *insertionKmerLeft ←*Insertion at location lo with base pair b and shift left
               *deletionKmerRight ←*Deletion at location lo and add base pair b at right extreme
               *deletionKmerLeft←* Deletion at location lo and add base pair b at left extreme
               Add all the Kmers formed above into an array
        **return** array of all the Kmers formed

**function** CREATESENTENCE(inputKmer) *⊳* Function to create a sentence for the given inputKmer which will be used for training
    *kmerArray ←*NEARBYSET(inputKmer)
    *sentenceArray←* []*⊳*Sentence with inputKmer at the center
    **for** kmer in first half of kmerArray **do**
          sentenceArray.append(kmer) sentenceArray.append(inputKmer)
    **for** kmer in second half of kmerArray **do**
          sentenceArray.append(kmer)
    *finalSentence ←*Join the words in sentenceArray using space to create a single string
    **return** finalSentence

**procedure** ENCODINGTRAINING
    *allKmers ←*Array containing all possible kmers
    *trainingData ←* []
    **for** kmer in allKmers **do** *⊳* Generate data for every kmer
          *kmerSentence←* CREATESENTENCE(kmer) trainingData.append(kmerSentence)
          *n*2*vEncodings←* Nucl2Vec Encodings trained on trainingData
     **return** n2vEncodings
~~~

mer length words composed of A,C,G and T. The training algorithm is described in Algorithm 1

A few random 4-mer encodings are represented in Fig 5 trained using the above method. We can see the proximity of 4-mers like AATG and AATC (edit distance = 1), CGAC and GACT (edit distance = 1). The 4-mers AAGG and GACC (edit distance = 3) are sufficiently far from each other. So are AAGG and CGAT (edit distance = 4). There are also a few anomalies present. Like AATG and AAGG (edit distance = 1) are not placed nearby while CGTT and GACC (edit distance = 4) are in close proximity of each other.

**Fig. 5.**
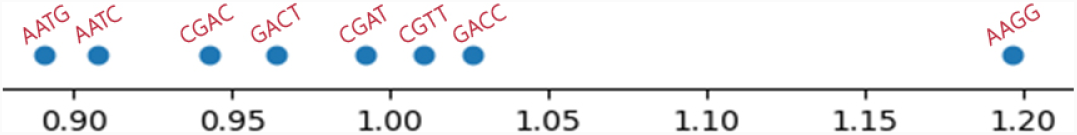
Comparing a few random encodings

Encoding k-mers into smaller dimensions provides a great boost in the computation speed. Yet there is also a clear disadvantage present in using these encodings due to the anomalies observed above. However if the probability of no correct match of any segment due to this encoding is, say *p*, in case of a single segment used, then it decreases to *p*p* for two segments used, *p*p*p* for three segments used and so on. In conclusion, the more number of segments we use for matching, the lesser chance there is of no segment matching at the correct position.

### B. K-Nearest Neighbors

KNN (K-Nearest Neighbors) algorithm is one of the simplest classification algorithms and it is one of the most used learning algorithms. KNN creates a tree from all available data cases and uses this tree to predict the classification of a new sample point based on some similarity measure. It is a non-parametric technique, which means that it does not make any assumptions on the underlying data distribution [15].

**Algorithm 2** Alignment

~~~
**function** ENCODEPMER(inputPmer, n2vEncoding, k) *⊳* Function to encode the inputPmer using n2vEncoding values
      *inputKmers ←* []
       **for** i from 0 to len(inputPmer)-k with jump of ’k’ **do**
                 inputKmers.append(inputPmers[i:i+k])
       *FinalEncoding←* []
       **for** kmer in inputKmers **do**
                  *enc←* Encoding of kmer in n2vEncoding
                  FinalEncoding.append(enc)
       **return** FinalEncoding

**function** PROCESSREFERENCE(reference, n2vEncoding, p, k) *⊳* Function to encode the reference DNA and create K NN Tree using n2vEncoding values
       *allPmers←*[]
       **for** i from 0 to len(reference)-p with jump of ’1’ **do**
                allPmers.append(reference[i:i+p])
       *allEncodings ←* []
       **for** pmer in allPmers **do**
                *enc←* ENCODEKMER(pmer, n2vEncoding, k) allEncodings.append(enc)
       *referenceTree ←*Create K NN Tree using allEncodings and their alignment (array index)
       **return** referenceTree

**procedure** FINALALIGNMENT(referenceDNA, readsArray, n2vEncoding, p, k, thresholdDis) *⊲* Values of ’p’ and ’k’ are defined experimentally with the constraint that ’p’ should be a multiple of ’k’
        *knnTree ←*PROCESSREFERENCE(referenceDNA, n2vEncoding, p, k)
        *alignment←* []
        **for** read in readsArray **do**
             *front←* ENCODEKMER(read[0:p], n2vEncoding, k)
             *back ←*ENCODEKMER(read[len(read)-p:len(read)], n2vEncoding, k)
             *center←* ENCODEKMER(read[len(read)/2-p:len(read)/2], n2vEncoding, k)
             *frontPos, backPos, centerPos ←* Best Match Alignment Positions from knnTree
             *frontDis, backDis, centerDis←* Best Match Distance Values from knnTree
             *optDis, optPos ←*Minimum(frontDis, backDis, centerDis) and the corresponding position
             **if** optDis <= thresholdDis **then**
                   alignment.append(optPos)
                **else**
                   alignment.append(-1)
         **return** alignment
~~~

In our model, the encoded reference k-mers from previous step are used as “training data” for KNN for preparing KNN tree while vector L2 distance is used as a measure of similarity, thus hypothesizing that similar k-mers will have similar encoding. While the algorithm is usually considered computationally heavy, with enough optimizations and pre-processing we can reduce the computation burden of the prediction process.

The algorithm is known to work well with large datasets, however it carries with it the curse of dimensionality, which means the data sparseness increases exponentially with the feature dimension. We therefore need to find a balance between the number of dimensions and the computation time required. This further reinstate the requirement of highly efficient feature encoding model.

### C. Complete Model

The proposed model takes an existing reference sequence, and a set of NGS reads(to be aligned) as input and outputs an integer value for each read representing the index at which the read aligns the best or -1 if the read does not align properly anywhere in the DNA. Algorithms 1, 2 and Fig 6 describe the steps executed to obtain the final output.

**Fig. 6.**
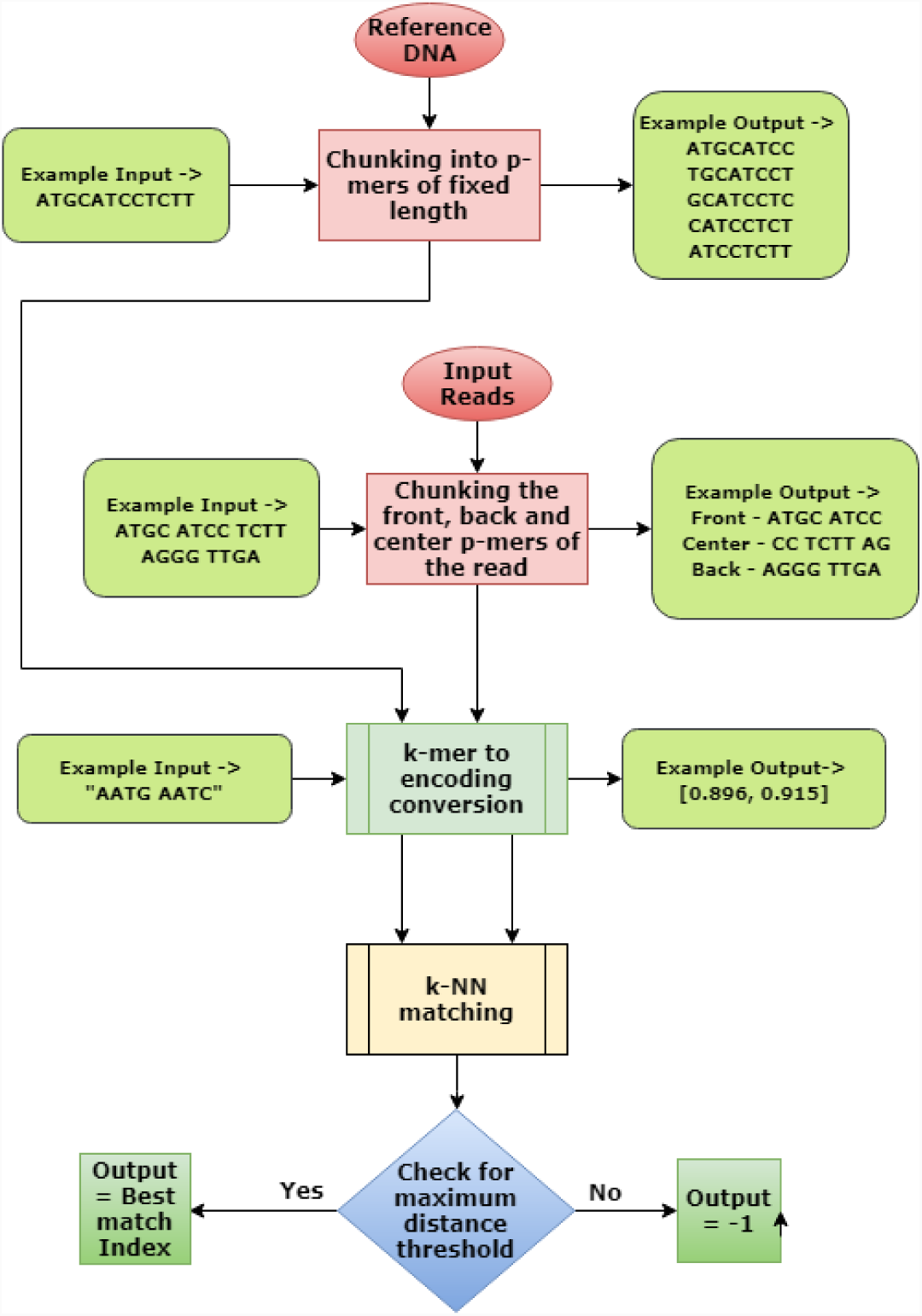
Complete Pipeline Flowchart

Step 1 : We first start with Word2Vec encodings of k-mers, which is learned separately. We use edit distance as previously described to represent proximity between two k-mer and learn the Word2Vec encodings and store them.

Step 2 : The pre-processing involves creating encodings of every p-mer present in the DNA (where p is a multiple of the encoding length k) and building a 1-Nearest Neighbor tree. For example, if the Word2Vec learned encodings for 4 length base pairs, then to encode a 16-mer, we first divide it into 4-mers and then encode them separately.

Step 3 : Once the encodings are learned and the preprocessing of the reference DNA is done, we proceed to align the reads. For each read, segments of length k from the front, back and center are taken and encoded. They are then passed through the 1-nearest neighbors tree created in the preprocessing for the best match. 1 nearest neighbour will output the match distance and predicted alignment location. Only matches which have the match distance less than some threshold are considered, for all others we output -1(unaligned).

### D. Dataset

To test our proposed model, we have used EColi Data generated form Illumina NGS with read length varies from 100-350 bps. The reference sequence is *CP012868.1 Escherichia coli str. K-12 substr. MG1655, complete genome*. The set of reads used for testing are mentioned in Table I

**TABLE I.** ACCURACY COMPARISON OF MODEL WITH BWA

| Name | Cat1<br>(%) | Cat2<br>(%) | Cat3<br>(%) | Cat4<br>(%) | Cat5<br>(%) | Cat6<br>(%) | Accuracy<br>(1+2+5) | Our Pred Time<br>(sec) | BWA-MEM Pred Time<br>(sec) |
| --- | --- | --- | --- | --- | --- | --- | --- | --- | --- |
| SRR2724094_1 | 93.57 | 0.15 | 0.62 | 0.95 | 3.66 | 1.20 | 97.39 | 19.002289 | 53 |
| SRR2724094_2 | 90.68 | 0.17 | 0.61 | 0.98 | 5.70 | 1.84 | 96.54 | 18.832064 | 61 |
| SRR2724099_1 | 91.55 | 0.16 | 0.62 | 0.67 | 4.86 | 2.13 | 96.57 | 51.007892 | 263 |
| SRR2724099_2 | 90.36 | 0.13 | 0.58 | 0.83 | 4.96 | 3.14 | 95.44 | 53.232324 | 211 |

## IV. Model Analysis

There are a number of hyper-parameters on which our model can be optimized. The analysis of each hyper-parameter has been described in the following subsections:

### A. Nucl2Vec Training Method

We trained our model for 3, 4, 5 and 6-mers with encoding dimension 1. We trained the model using both the traditionally used heuristic based training methods and the thorough training methods that we propose. It is clear from Fig 7 that our training method (thorough training without using any heuristics) consistently performs better than the traditional Word2Vec training method.

**Fig. 7.**
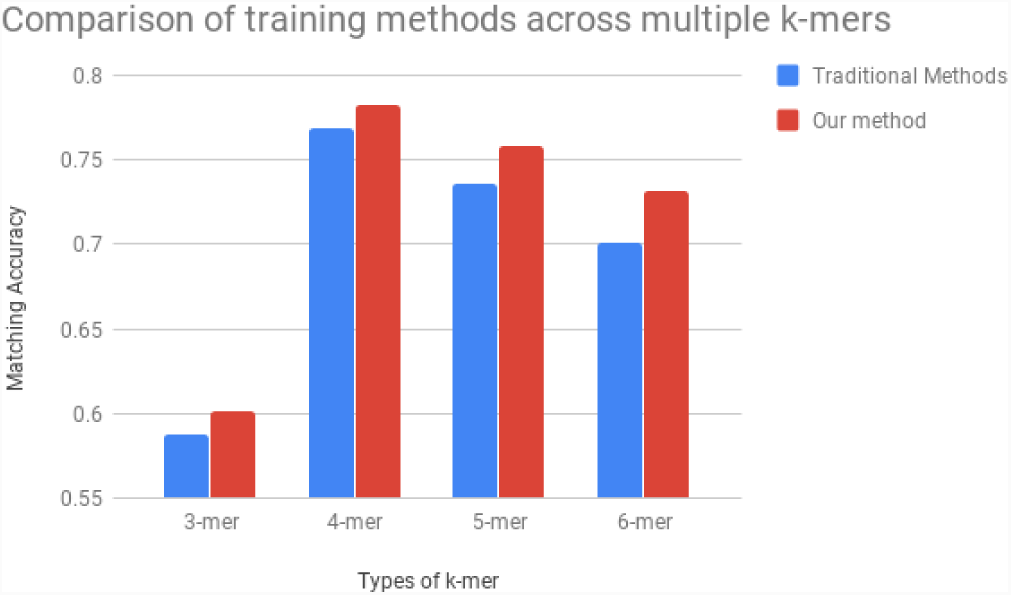
Accuracy Comparison for traditional Word2Vec training methods vs our training method

### B. Encoding Length

The prediction time for a n-dimensional 1-nearest neighbors matching is proportional to ’n’. Thus, converting bigger k-mers into single encodings will provide us with more portion of the read to match without any extra computation. However, the longer sequences we try to encode into 1-dimension, the more error prone these encodings become. So there needs to be a balance among the two. [fig9]

From Fig 8, we can see that encoding 4-mers seems to be the correct choice for the best accuracy. Also from Fig 9, we can see the variation in computation time between encoding in 1- dimension vs encoding in 2-dimension and since the difference in accuracy between the two is not significant, we should prefer encoding in 1-dimension for better prediction speed.

**Fig. 8.**
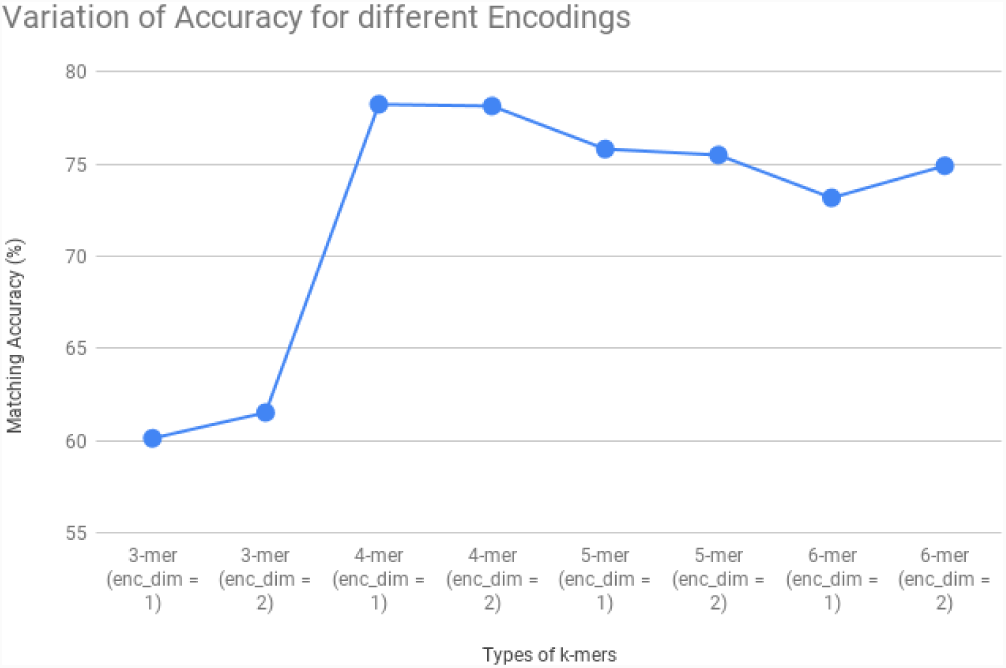
Variation of Matching Accuracy for different Encodings

**Fig. 9.**
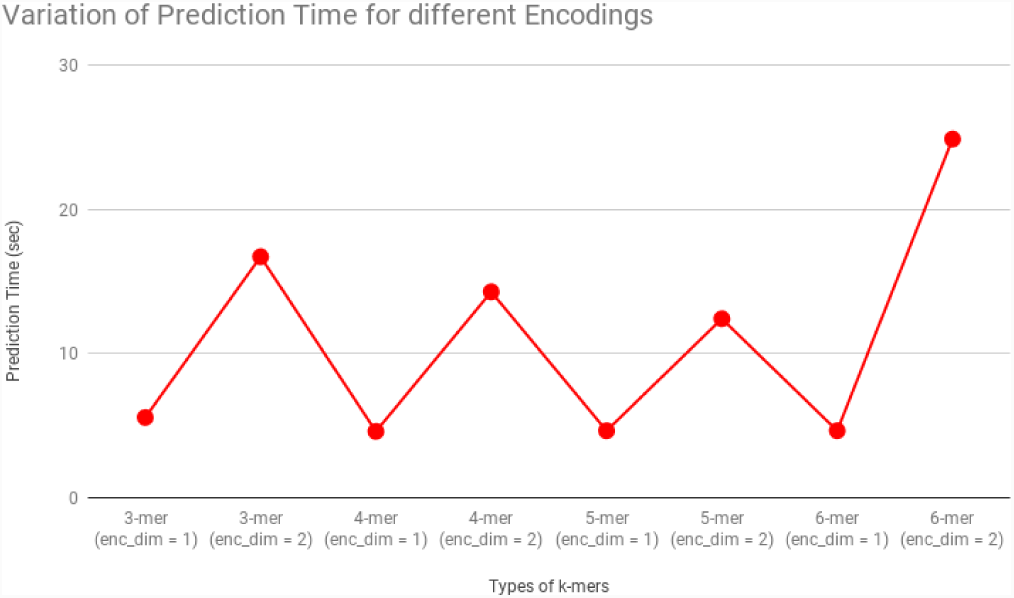
Variation of Prediction Time for different Encodings

### C. Length of the matching Segment

Now that we have experimented with the k-mer size and the encoding lengths and discovered that the best encodings are 4-mers with encoding dimension = 1, we will use these values for all future analysis.

We experimented with different length segments to match and plotted the accuracy and prediction time against the length of segment used. From Fig 10, it can be seen that the accuracy reaches a certain threshold at segment length = 16 base pairs, after which it does not increase but instead decreases (this may happen due to the curse of dimensionality). Also we can see from Fig 11 that the prediction time increases exponentially as expected.

**Fig. 10.**
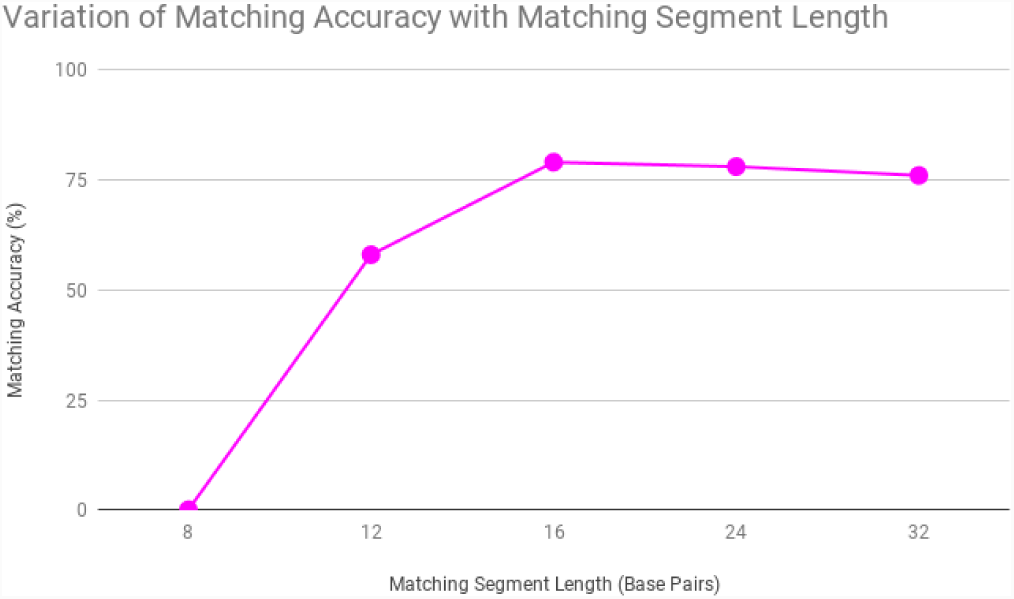
Variation of Matching Accuracy with Matching Segment Length

**Fig. 11.**
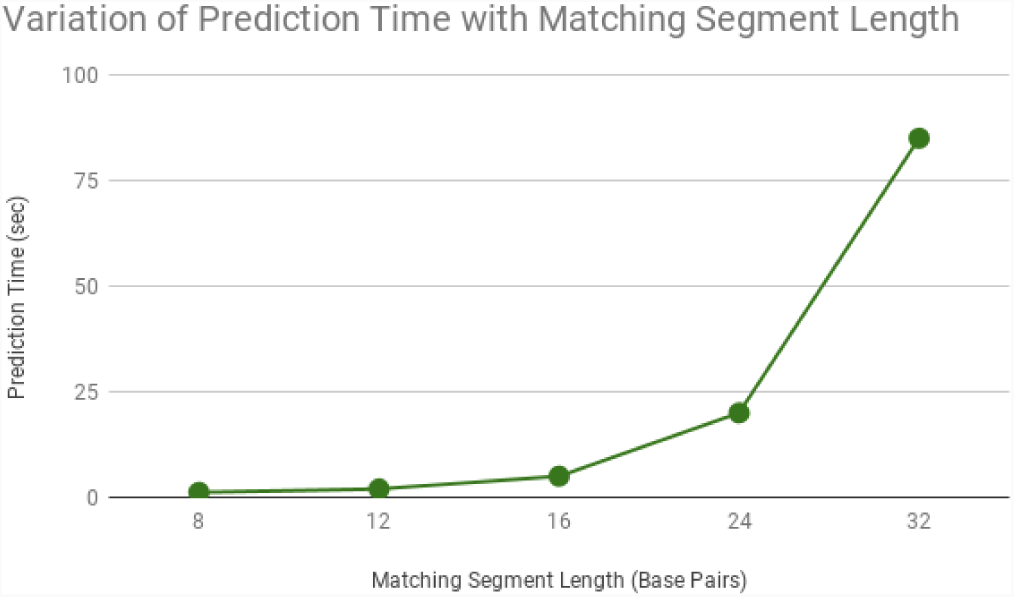
Variation of Prediction Time with Matching Segment Length

### D. Number of Matching Segments

Since we are using 16-mer segments for alignment, we have to determine how many and from where these segments should be taken. The possibilities include the front segment, the back segment, the left of the center and the right of the center. We experimented with the combinations of these and the accuracies obtained and the prediction time were plotted. From Fig 13, it can be seen that the prediction time increases linearly with the number of pieces used and we can also see from Fig 12 than the accuracy starts saturating as we increase the number of segments used. There is still a significant increase in accuracy even between using 3 segments and 4 segments, thus we will use 4 segments for final results.

**Fig. 12.**
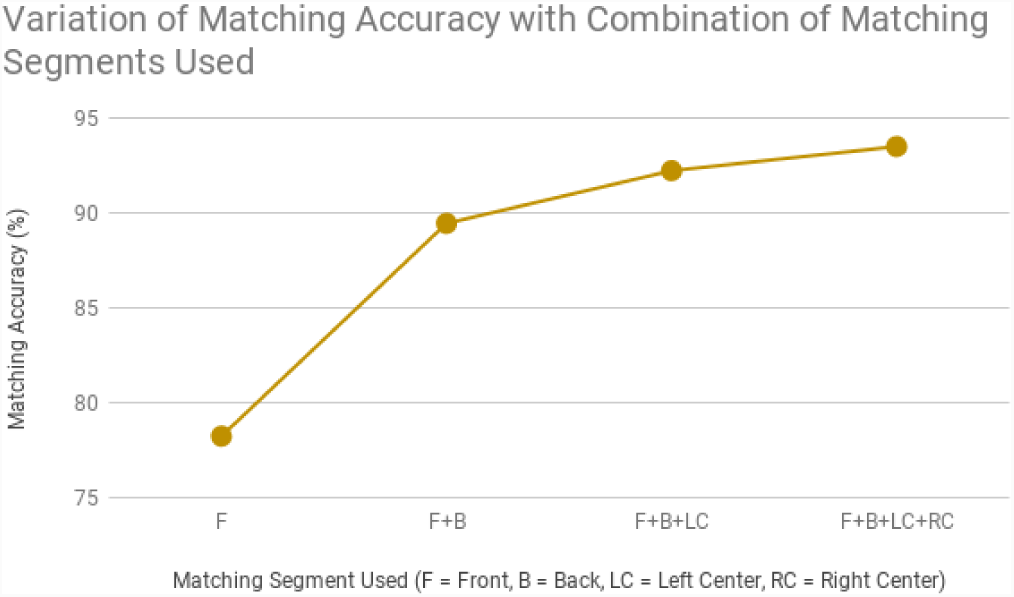
Variation of Matching Accuracy. F = Front, B = Back, LC Left Center, RC = Right Center

**Fig. 13.**
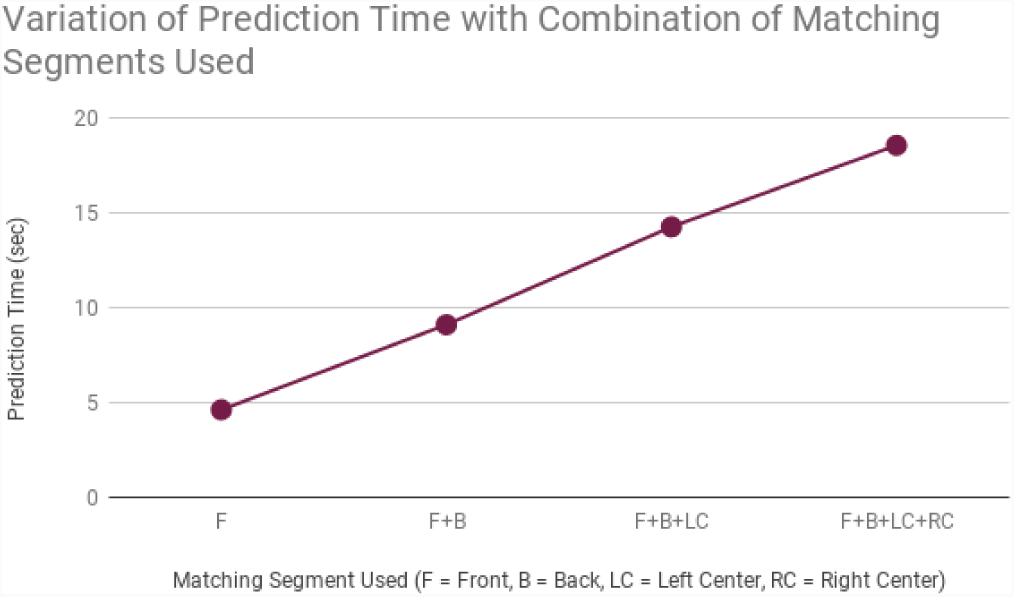
Variation of Prediction Time. F = Front, B = Back, LC Left Center, RC = Right Center

### E. Distance Threshold

Our model will always predict an alignment location corresponding to every read. However, to reject substandard matches, we require some threshold on maximum distance of aligned segment. To calculate this threshold, we randomly picked 200 reads which had a matching index and 200 reads which were not supposed to match at any index. We plotted the matching distance output from our algorithm against the number of reads. From Fig 14, we can see that the red dots (representing no match) have a higher frequency at larger distances and the blue dots (representing a match) have higher frequency at lower distance values.

**Fig. 14.**
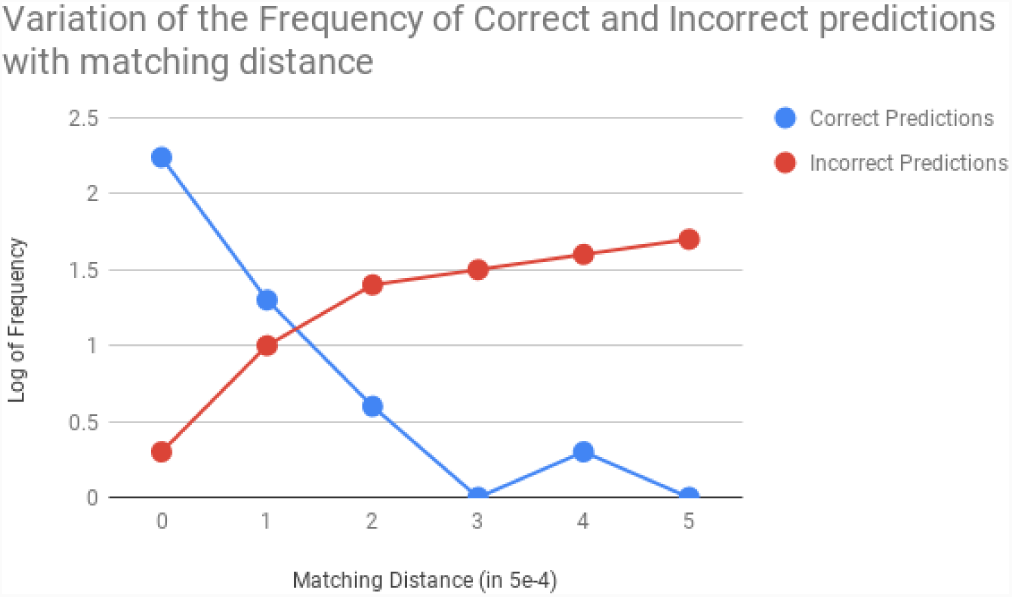
Distribution of Matching Distance with the log of Frequency of correct and incorrect matches.

The average distance of all the reads which should match was 0.000111579, while the average of no matches was 0.002353325. So we placed a threshold at 0.0002.

## V. Verification of Predicted Results

In the previous section we have compared the accuracy of our model against BWA-mem results. To analyze the mismatch instances we have used Needlemen Wunsch algorithm to calculate alignment score. We observed cases where our predictions are penalised despite better NW scores.

1. Where our results do not match BWA results, but NW scores of both alignments are comparable. This might be due to the possibility of a read aligning at multiple locations in the reference.
2. Where BWA predicted the read to be unaligned, but our model predicted an alignment with acceptable NW score.
3. Our predicted alignment is offset from the correct alignment position and our NW score is low as compared to score at BWA predicted location. This may be due to two reasons: one is due to a few inaccurate Word2Vec encoding as we have discussed earlier and other reason may be different interpretation of best local alignment by CIGAR(BWA) and by KNN distance (see fig 15).

The errors from third case can be corrected by doing NW score check in the local neighborhood of our predicted location. Since NW takes considerable amount of computation, this has not been included in our final pipeline. However the same is analyzed in a little detail below. We need to find how much extra string from both sides of the predicted index can be considered reasonable to consider while finding the best CIGAR

**Fig. 15.**
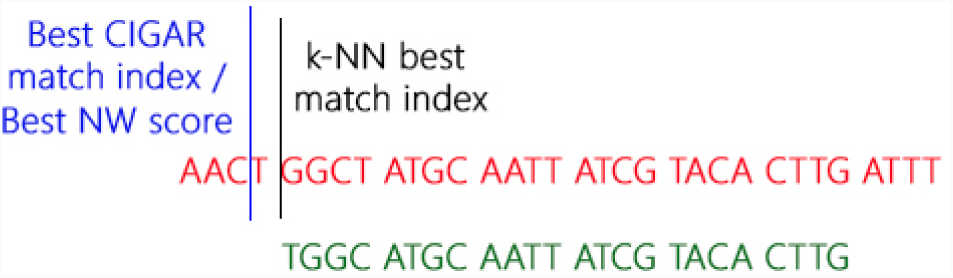
Local Alignment using NW score

So we plotted the distance between the actual index and the predicted index for reads that our algorithm was not able to match against its frequency (Fig 16). We have not included distance greater than 80 base pairs, since that means the alignment was done at a completely different place and using NW algorithm locally will not help eliminate those errors.

**Fig. 16.**
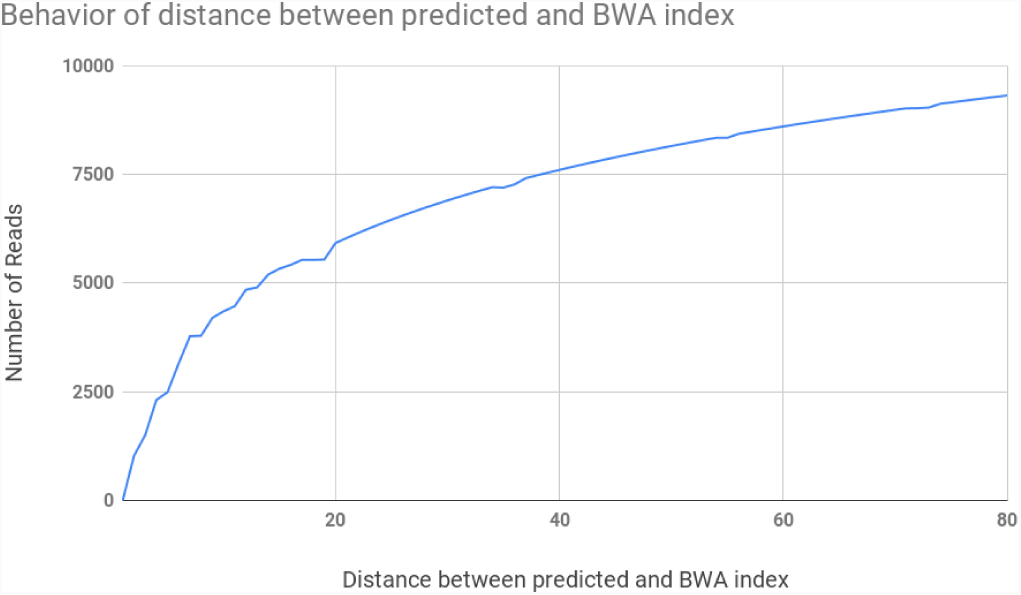
Variation of Number of Reads with distance between predicted and BWA index less than or equal to the given value

We can see from Fig 16 that more than 50% of all the errors can be eliminated by just doing a local alignment at a distance of 20.

### A. Final Analysis

To compare the final results, 4-mers encoded into encoding dimension = 1 were used. Each segment of read used for matching was 16 base pairs long, and 4 such segments were used (Front, Back, Left Center and Right Center).

The final results were divided into six categories –

1. Category 1 : Predicted Labels same as BWA labels.
2. Category 2 : BWA Labels say no match but predicted label contains match with acceptable NW score.
3. Category 3 : BWA Label and Predicted Label differ, and the predicted label NW score is low.
4. Category 4 : BWA Labels say no match and predicted label contains match with a low NW score.
5. Category 5 : BWA Label and Predicted Label differ, however their NW scores are comparable.
6. Category 6 : Predicted Label is -1, but there was an alignment present in the BWA label.

The final accuracy comparisons and speed analysis against BWA-MEM are done in Table I. Clearly Category 1, 2 and 5 comprise the correct prediction percentage and the rest three categories amount to error percentage.

## VI. Challenges and Future Work

The aim of the paper is to provide a way of generating features out of k-mers that represent more information than the traditional one-hot vector encodings. The Nucl2Vec embeddings proved to be a good encoding method for the same. However, the encodings used by this paper were trained based on purely mathematical intuitions only. Substitutions of certain kind might have more biological preference than others and thus this information can be used to determine k-mers proximity while training Nucl2Vec encodings. Adding such biological insights into the encoding can increase the accuracy further.

Following the encodings of the base pairs, we used k-nn algorithm for alignment. Different algorithm can also be used and experimented with using the same encodings for faster and better alignment. The code used in this paper along with its documentation is readily available for reproducibility here.

